# Quantitative multiplexed ChIP reveals global alterations that shape promoter bivalency in ground state embryonic stem cells

**DOI:** 10.1101/557082

**Authors:** Banushree Kumar, Simon J Elsässer

**Author notes:** **Corresponding Author**: Simon J Elsässer.

## Abstract

To understand the epigenetic foundation of naïve pluripotent cells, we implement a quantitative multiplexed ChIP-Seq method (MINUTE-ChIP) comparing mouse ESC grown in 2i versus Serum conditions. Combined with quantitative western blot and mass spectrometry, we find compelling evidence for a broad H3K27me3 hypermethylation of the genome, concomitant with the widespread loss of DNA CpG methylation. We show that opposing action of EZH2 and JMJD3/UTX shape the H3K27me3 landscape, with almost all bivalent promoters stably retaining high H3K27me3 levels in 2i. In parallel, we show a mechanistically uncoupled, global decrease of H3K4me3 and strong reduction at bivalent promoters, suggesting that low H3K4me3 and high H3K27me3 levels at bivalent promoters safeguard naïve pluripotency.

## INTRODUCTION

The mouse pluripotent ground state is attributed to naïve epiblast cells of the inner cell mass and can be recapitulated *ex vivo* through inhibition of GSK-3b and MAPK/ERK signaling in mouse ESC (mESC) in serum-free conditions containing leukaemia inhibitory factor (LIF) and two respective inhibitors (CHIR99021 and PD0325901, referred to as “2i”) ^1^. Traditional serum-based culture in the presence of LIF (hereafter “Serum”) maintain a more metastable ES cell phenotype. While rapidly interconvertible through change in media condition, significant differences in signaling, metabolism, transcriptional regulation, epigenomes and nuclear organization between the two states have been reported ^1–7^. Given that metabolic changes often induce global shifts in histone modification patterns, we reasoned that revisiting existing paradigms on epigenomic changes between the two states in a quantitative manner would be warranted. We implemented a multiplexed ChIP-Seq method to compare epigenetic profiles between 2i and Serum conditions. We focused on two histone modifications, H3K4me3 and H3K27me3 known to constitute developmentally important bivalent chromatin domains ^8^. Prior studies agree on a model where bivalent domains are resolved in ground state pluripotent cell into H3K4me3-only domains through the loss of H3K27me3 ^2–5, 9, 10^.

## RESULTS AND DISCUSSION

Barcoding-first ChIP techniques have gained popularity because of the combined benefit of multiplexing and quantitative readout ^11–14^. Pooling barcoded samples greatly increases throughput without complicated automation, while effectively removing technical variability between samples. Challenges remain for fragmentation and ligation of crude chromatin in a manner that maximizes barcoding efficiency while avoiding any technical variability or sample loss that could confound the subsequent quantitative measurement. Mint-ChIP, developed by the Bernstein lab ^12^, provides a formidable solution for these problems with a streamlined one-pot chromatin barcoding and a post-ChIP linear amplification that requires only one adaptor per chromatin fragment. In short, native chromatin is fragmented using Micrococcal nuclease, and subsequently blunted and ligated to double-stranded DNA adaptors that include a T7 RNA polymerase promoter and a sample barcode sequence. Finally, samples are combined and subsequent ChIP reactions are performed with the pooled samples. ChIP material is prepared into an Illumina-compatible library using linear amplification by virtue of T7 RNA polymerase, reverse transcription and a low-cycle library PCR amplification ^12^.

Here, we introduce unique molecule (UMI) counting and paired-end mapping of the chromatin fragments to this method, which we then termed MINUTE-ChIP for multiplexed indexed unique molecule T7 amplification end-to-end sequencing (Figure 1a, Supplementary Table 1). Double-stranded DNA adaptors were used to barcode chromatin in a blunt-end ligation reaction. As in the original Mint-ChIP design, adaptors carried a partial SBS3 for Illumina sequencing flanked by a T7 RNA Polymerase for linear amplification. Between the SBS3 sequence and a 8bp sample barcode at the 3’ end, a 6bp randomized sequence was introduced, serving as a unique molecular identifier (UMI). Using UMI information in addition to the ligated genomic sequence greatly increases the confidence in calling amplification duplicates and improves the quantitative representation of repetitive sequencing (Supplementary Figure 1a). Pooling a human HeLa cell sample with the mouse ESC samples, adaptor-crosscomination was assessed to be very low, with a maximum of 1.5% mapping to the respective other species. (Supplementary Figure 1b). For paired-end mapping, we modify the linear amplification strategy introduced in Mint-ChIP by priming the cDNA synthesis from a 3’ RNA adaptor, thus maintaining the original genomic fragment length, retrieving a typical mononucleosomal fragment length for histone ChIP (Supplementary Figure 1c, Supplementary Table 1). As previously reported ^12^, barcode representation in the pool may vary amongst samples, even at precisely matched adaptor DNA concentration. For accurate quantification, barcode representation after the ChIP has to be related back to the corresponding quantities in the input pool. A common issue of indexing-first ChIP protocols is that adaptors added in excess are carried over into the input pool. Amplification products from free adaptors, adaptor dimers or other side reactions contribute contaminating sequences to an extent that sequencing of the input pool becomes unfeasible. Thus we have optimized the stoichiometry of adaptors to chromatin to enable sequencing of the input. Quantification against input representation is robust, as exemplified by a H3 ChIP from a pool containing two series of biological triplicate samples with varying input representation; all replicates lie within 10% variance (Figure 1b, Supplementary Figure 2, Supplementary Table 2).

**Figure 1.**
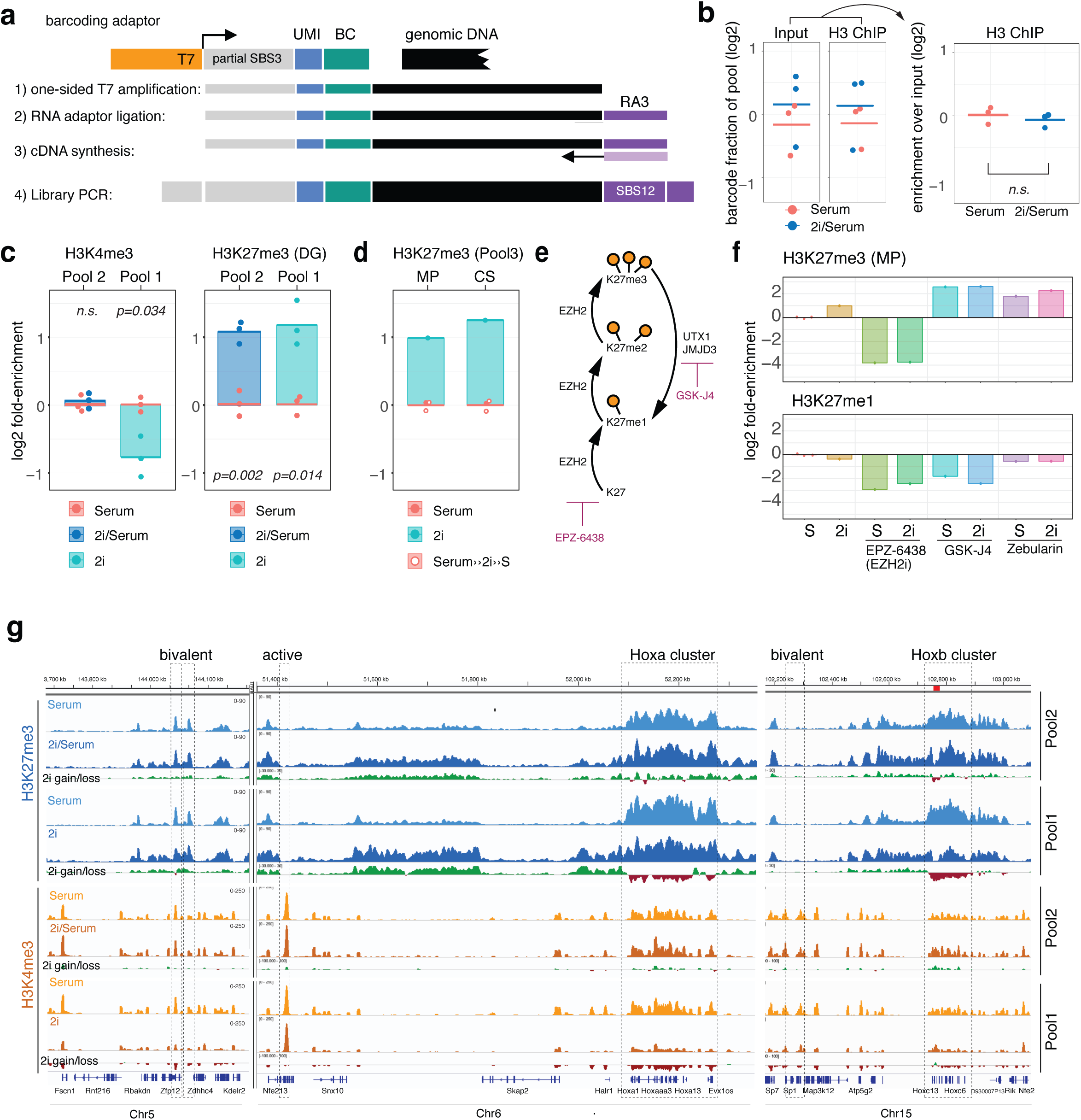
MINUTE-ChIP reveals high H3K27me3 and low H3K4me3 levels in naïve mouse ESC. **a** Schematic of MINUTE adaptor design for barcoding chromatin. One-sided ligation of adaptor comprising T7 RNA Polymerase promoter (T7), random 6bp sequence (UMI) and 8bp barcode (BC) is sufficient for subsequent linear amplification. cDNA is primed from a ligated RNA adaptor (RA3). SBS3 and SBS12 designates standard Illumina read1 and read2 sequencing primers. **b** Example of quantification. H3 ChIP was performed on a pool (Pool2) of two condition with three biological replicates each. Barcode representation before and after ChIP (left), and resulting quantitative comparison (right). **C** Quantitative comparison of H3K4me3 and H3K27me3 levels in biological triplicates of Rw4 mESC grown in Serum vs 2i condition (Pool1), Serum vs 2i/Serum condition (Pool2). H3K27me3 antibody was from Diagenode (DG). **d** Quantitative comparison of H3K27me3 (Pool3) in RW4 mESC grown in Serum, switched to 2i for three passages (2i) and back to Serum for three passages (Serum>>2i>S). H3K27me3 antibodies were from Millipore (MP) and Cell Signaling (CS). **e** Schematic of enzymes involved in adding and removing H3K27 methylation, and specific inhibitors used in Pool3. **f** Quantitative comparison of H3K27me3 and H3K27me1 in Serum or 2i conditions, with specific inhibitors. See Supplementary Figure 3 for full dataset description. **g** Example H3K27me3 and H3K4me3 tracks from Pool1 and Pool2. Scaling reflects quantitative differences within each pool. Biological replicates were combined into single tracks. An additional track is showing gain and losses for each pool and modification.

### High levels of H3K27me3 and low levels of H3K4me3 characterize 2i ground state

Here, we performed three independent MINUTE-ChIP experiments, “Pool 1” comparing Serum and serum-free 2i conditions, “Pool 2” comparing Serum condition without or with 2i “2i/Serum”, and “Pool 3” including conditions with inhibitors against the H3K27me3 methyltransferase EZH2 and demethylases JMJD3/UTX (Supplementary Figure 3). Unexpectedly, quantitation of these three MINUTE-ChIP experiments, using three different H3K27me3 antibodies, (for details and primary data see Supplementary Table 2) all reported two-fold higher levels of H3K27me3 in the presence of 2i (Figure 1c-f). We used both mass spectrometry and quantitative western blot to validate our finding. Indeed, both measurements suggested an increase in total H3K27me3 (Figure 2c, 2d; 1.5-fold and 1.7-fold, respectively) (Supplementary Figure 4). These results are surprising particularly on the background of the widely accepted model that H3K27me3 is lost from Polycomb targets such as bivalent domains and Hox gene clusters ^2–5, 9, 10, 15^. However, a recent study also reassessed H3K27me3 levels in a quantitative manner and reports a similar global increase, concomitant with a gain in PRC2 components ^16^.

**Figure 2.**
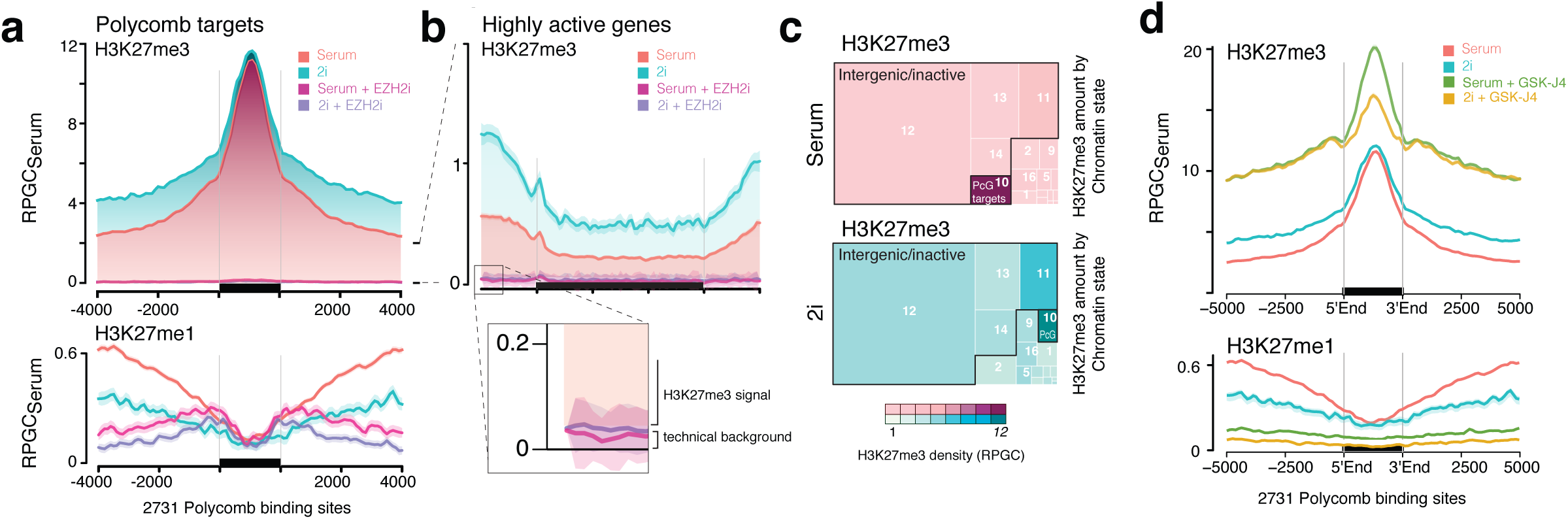
Widespread H3K27me3 methylation in pluripotent mESC is exacerbated in 2i. **a** Average profile of H3K27me3 and H3K27me1 levels at 2731 PcG target sites defined by EZH2 or Ring1b binding, in Serum and 2i condition, as well as after treatment with EZH2 inhibitor EPZ-6438 (EZH2i) for 7 days. Y axis is normalized to a global mean coverage density of 1 (reads per genome coverage, RPGC) in Serum. See Supplementary 7 for corresponding heatmaps. **b** Average profile at 1670 highly transcribed regions exhibiting lowest genome-wide H3K27me3 levels. Y axis is normalized to RPGC in Serum. Inset shows magnification of H3K27me3 signal in EZH2i treated conditions. **c** Treemap of H3K27me3 distribution across 17 functional chromatin (ChromHMM) states (listed in Supplementary Figure 6). Area is true to total proportions of H3K27me3. Color gradient shows relative density (total H3K27me3 normalized to proportion of each state of total genome size). **d** Average profile of H3K27me3 and H3K27me1 levels at PcG target, in Serum and 2i condition, as well as after treatment with demethylase inhibitor GSK-J4. Y axis is normalized to RPGC in Serum. See Supplementary Figure 7 for corresponding heatmaps.

Surprisingly, we also found significant perturbation of the H3K4me3 levels, but in a different pattern: Global H3K4me3 levels were 1.8-fold decreased in 2i but not 2i/Serum (Figure 1c), also evident from quantitative western blot (Supplementary Figure 4), representing another global difference that previously escaped observation. This suggests that independent mechanism regulate H3K27me3 and H3K4me3 levels. The former is linked to the action of 2i inhibitors, whereas H3K4me3 levels are insensitive to 2i inhibitors but respond to removal of serum.

Next, we wondered how these significant global perturbations of histone modifications would affect local chromatin environments. A first inspection suggested a broad gain in H3K27me3 across gene-poor regions and unchanged enrichments at peaks corresponding to bivalent promoters (Figure 1g). As an exception to this, Hox clusters showed a variable reduction in H3K27me3, albeit much less pronounced than previously described ^3, 16^. However a region in the Hoxc cluster (Hoxc12-c13), shown to be transcriptionally activated in 2i, was also fully depleted of H3K27me3 in our experiment (Supplementary Fig 5). Further, UMI deduplication allowed us to also quantitatively compare relative enrichment of histone modifications between the annotated genome and repetitive regions. H3K27me3 was relatively depleted from major and minor satellite repeats in Serum but increased in 2i albeit below the genome average (Supplementary Figure 6).

The H3K4me3 landscape appeared unaltered in 2i/Serum. In 2i however, most H3K4me3 peaks appeared reduced, most prominently at bivalent promoters (Figure 1g). Notably, it has been previously shown that bivalent promoters show somewhat reduced H3K4me3 in 2i condition, and that this is a rapid response within 24 hours of switching Serum-grown cells into 2i condition ^5^. This lead us to hypothesize that high H3K27me3 and low H3K4me3 are two mechanistically unlinked characteristics of the ground state, and that the 2i/Serum conditions allows to stabilize a normally transient intermediate between Serum and 2i conditions.

### Polycomb binding sites maintain high levels of H3K27me3 in 2i

The enzymes responsible for methylation and demethylation of histone H3K27 are well established: EZH2 is the enzymatic subunit of PRC2 that catalyzes all three H3K27 methylation states; Two demethylases, UTX and JMJD3, are known to convert H3K27me3 to H3K27me1 (Figure 1e). Thus, we were particularly interested how the H3K27 methylation landscape of ground state pluripotency is sculpted by these two activities and profiled H3K27me3 and H3K27me1 (Pool 3). We focussed on well-described polycomb (PcG) binding sites combining EZH2 and Ring1b published peak sets (Figure 2a, Supplementary Figure 7. Notably, average H3K27me3 levels at the peak were unchanged in 2i, whereas neighbouring regions gained H3K27me3, concomitant with a loss of H3K27me1 (Figure 2a). In fact, H3K27me3 was gained roughly two-fold, genome wide across most genomic contexts with the notable exception of several transposable elements, e.g. IAP ERVs (Supplementary Figures 6, 8). Such global shift confounds traditional ChIP-Seq analyses which are normalized based on the assumption of a stable technical background (Supplementary Figure 9).

### Pervasive H3K27me3 modification is a hallmark of mouse ESC and further exacerbated in 2i conditions

In any ChIP experiment, a certain level of technical background is expected from unspecifically retained DNA contamination. To determine the technical background and dynamic range of our MINUTE-ChIP experiments, we evaluated H3K27me3 ChIP in the presence of an EZH2 inhibitor EPZ-6438 ^17^, hereafter (EZH2i). Notably, barcodes for the EZH2i treated samples were depleted 14 to 30-fold from the pool after ChIP. Treatment with GSK-J4 an inhibitor of UTX and JMJD3 on the other hand, resulted in a 6-fold increase (Figure 1f, Supplementary Figure 10). Thus, technical background is extremely low, contributing roughly 1% of the maximal signal. Even at regions known to be devoid of H3K27me3, such as highly transcribed genes, this technical background is substantially below the measured signal (Figure 2b). This observation has important implications to interpreting our, or any, H3K27me3 ChIP profiles: H3K27me3 modifications are omnipresent on nucleosomes at substantial densities across the genome. Despite the high local density (up to 100% as suggested by calibrated ChIP ^18^), only a small fraction (3.6% in Serum and 2.0% in 2i) of H3K27me3-modified nucleosomes reside in PcG target regions (Figure 2c). Mechanistically this implies, for example, that any effector proteins downstream of H3K27me3 must be exquisitely sensitive to the density of H3K27me3 present on the chromatin fiber to selectively bind PcG target regions and not be titrated away by bulk H3K27me3 in the rest of the genome.

### Gain in H3K27me3 follows the loss of CpG methylation

Turning back to our question of how ground state pluripotency invokes a broad increase in H3K27me3 at non-PcG targets, we noted similarities to a previously observed retargeting of PRC2 in mESC devoid of CpGme ^7, 19^. A known feature of ground state ESC is a strong decrease in 5-methyl cytosine methylation at CpG dinucleotides (CpGme) ^5–7^. Indeed, H3K27me3 is predominantly gained in regions that switch from high to low CpGme levels in 2i (Supplementary Figure 11). ERV families which are known to retain high levels of CpGme during germline reprogramming and also in 2i ^6, 7, 20^, continue to be depleted of H3K27me3 (Supplementary Figure 6). Regions with constitutively low CpGme, such as the above-mentioned PcG target sites, stably maintain their high H3K27me3 levels. Depleting CpGme with Zebularin in Serum or 2i condition very broadly increased H3K27me3, thus in a less specific way then the Serum-2i transition. This can be explained by the fact that cells in 2i conditions are not completely devoid of DNA methylation. While our results corroborate the hypothesis that unmethylated CpGs are a major determinant of the H3K27me3 landscape, the ground state epigenome certainly underlies are more complex regulation since Zebularin treatment e.g. did not recapitulate the specific loss of H3K27me3 at Hoxc cluster. This is in line with the large body of evidence that DNMT knockout or otherwise reduced CpG levels per se do not induce the pluripotent ground state ^5–7, 19^. Nevertheless, it may be speculated that CpG demethylation is necessary albeit not sufficient for H3K27me3 hypermethylation and naive state. In fact, a recent bioRxiv preprint suggests that restoration of Serum-like DNA methylation in the naive state also reverts back the H3K27me3 landscape, albeit the cell maintains a naive transcriptome and phenotype ^21^. Vice versa, H3K27me3-deficient EED^-/-^ mESC have been shown to exhibit increased CpG methylation in 2i ^22^, while also maintaining the naive transcriptome ^5^.

### Active demethylation limits H3K27me3 levels genome-wide

GSK-4 treatment further revealed that demethylases UTX and JMJD3 limit H3K27me3 levels both at PcG targets and genome-wide (Figure 2d, 1f, Supplementary Figure 12). Analysis of the gain/loss of H3K27me3 according to initial H3K27me3 quantiles in serum revealed that GSK-J4 favors accumulation of H3K27me3 in regions exhibiting low to very low H3K27me3 while affecting H3K27me high regions to a lesser extent (Supplementary Figure 12a). This would fit to a model in which UTX/JMJD3 act relatively unspecific on H3K27me3 scattered across the genome, limiting it’s abundance predominantly non-Polycomb targets. Such model is also in agreement with genome-wide maps of UTX/JMJD3 that found a relatively flat distribution with weak enrichment at active gene promoters rather than Polycomb targets ^23, 24^. Interestingly, GSK-J4 treatment assimilated H3K27me3 genome-wide distribution in serum and 2i almost completely (Supplementary Figure 12b). Thus, it remains to be determined if and how their localization and/or activity is modulated by the Serum-to-2i transition. In summary, PRC2 and UTX/JMJD3 jointly balance H3K27me3 level in both serum and 2i conditions.

### Maintenance of H3K27me3 and loss of H3K4me3 at bivalent promoters

Compared to Serum condition, the 2i ground state is characterized by low expression of particular lineage-specific transcription factors that are thought to prime differentiation. The apparent loss of H3K27me3 at the bivalent promoters of these genes has been difficult to reconcile with a repressive role of the modification, and the fact the the target genes were maintained silent ^3^. Overlapping largely with the PcG target sites discussed above, we find that H3K27me3 is quantitatively maintained at the ∼2000 bivalent promoters (Figure 3a-c, Supplementary Figure 13). While individual datasets showed more variability in H3K27me3 peaks, 171 promoters lost and 126 gained H3K27me3 consistently more than 1.5-fold across our three independent MINUTE-ChIP experiments (Figure 3d, Supplementary Figure 14). Those gaining H3K27me3 were strongly downregulated. Those losing H3K27me3 included Hox clusters and regions previously shown to interact with Hox clusters ^4^, and showed a variable degree of upregulation (Supplementary Figure 15). Thus, our quantitative data reconciles apparent discrepancies arising from non-quantitative data and agrees with the common notion that H3K27me3 is anticorrelated with transcriptional activity.

**Figure 3.**
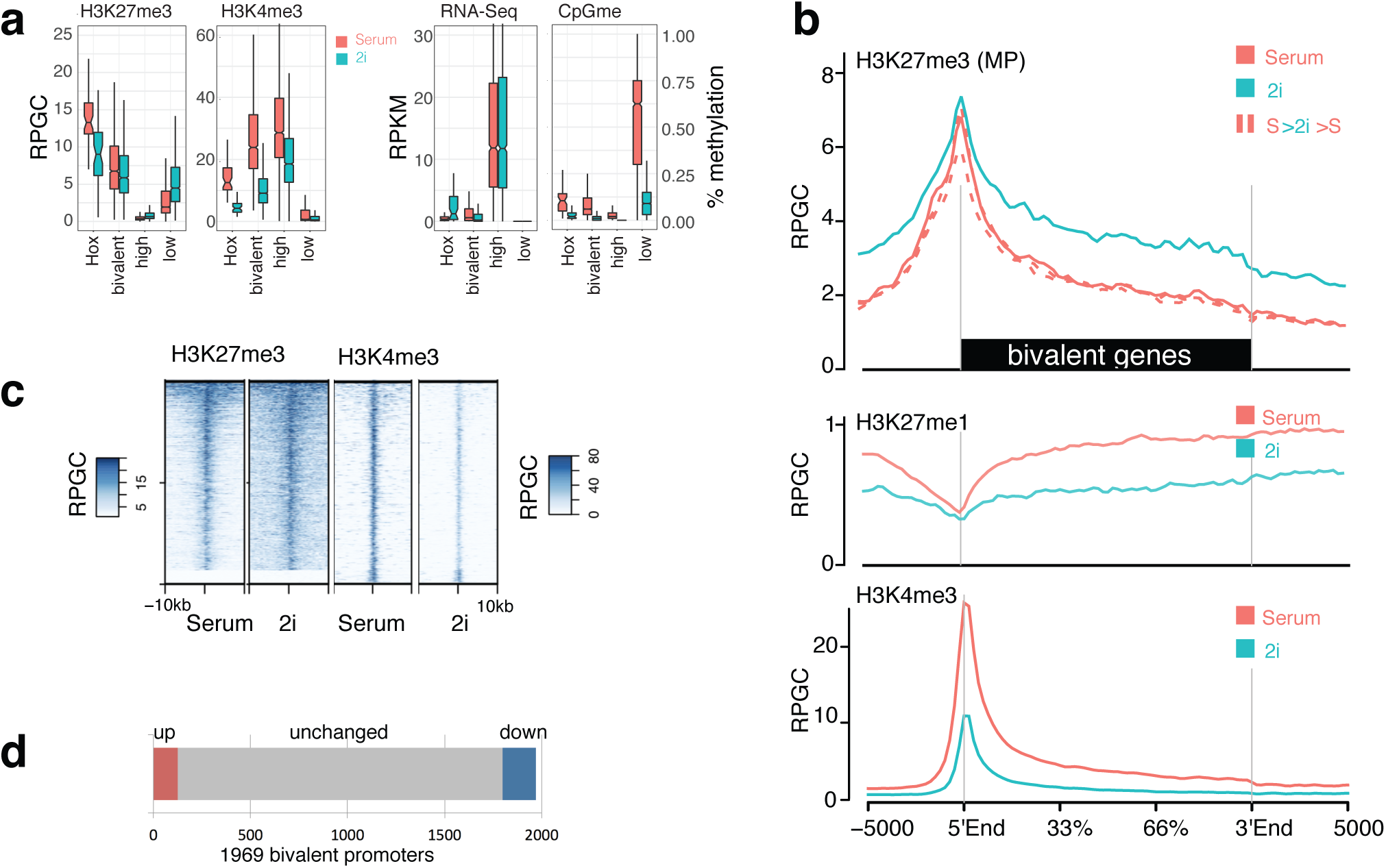
Bivalency safeguarding developmental genes in ground state pluripotency is characterized by low H3K4me3 and high H3K27me3. **a** Relative levels of H3K27me3 and H3K4me3 at bivalent, non-transcribed (low) and highly active (high) promoters, as well as Hox genes (Hox). Y axis is normalized to RPGC in Serum. See Supplementary Figure 13b for quantification across a more comprehensive set of functional regions. RNA-Seq expression levels and CpG methylation levels at the same promoters (right). **b** Average profiles of H3K27me3, H3K27me1 and H3K4me3 across 1969 bivalent genes. Additional H3K27me3 antibodies are shown in Supplementary Figure 13a. **c** Heatmap across promoters of the bivalent genes as in a). **d** Number of bivalent promoters showing 1.5-fold or higher change between 2i and Serum, consistent across the three different H3K27me3 ChIPs. The most up/ down regulated genes are shown individually Supplementary Figure 14. RNA-Seq expression levels of the up/down regulated genes is shown in Supplementary Figure 15.

The global reduction of H3K4me3 (Figure 1c) in 2i condition manifested in reduced peak sizes across all genes, but most strikingly at Hox genes and bivalent promoters. While active genes on average lost ∼35% of H3K4me3, bivalent promoters showed a more than 2-fold decrease in H3K4me3 (Figure 3a-c). This is in line with prior observations that H3K4m3 levels are reduced in 2i conditions with relative to other promoters ^5^. While loss of H3K4me3 has no apparent effect on the transcriptional output of active genes, the more dramatic loss at bivalent genes, in the presence of similar H3K27me3 levels, may account for the small but consistent decrease in transcription from bivalent genes (Figure 3a, Supplementary Figure 15).

While necessary for survival in the absence of 2i, serum primes mESC for differentiation. Our data suggests that high H3K4me3 at bivalent genes is a molecular manifestation of the serum-primed state. Indeed, a recent report suggests that H3K4me3 specifies the robust and timely induction of bivalent promoters during differentiation ^25^. Given that H3K27me3 is *per se* dispensable for ground state pluripotency ^5, 16^, we hypothesize that low H3K4me3 together with high H3K27me3 levels at bivalent promoters act to safeguard the ground state of pluripotency. Our study highlights the importance of performing quantitative ChIP experiments by uncovering unexpected global shifts in histone modifications that have confounded prior non-quantitative studies.

## METHODS

### Cell Culture

Rw4 murine embryonic stem cells (mESCs) were cultured feeder-free in 0.1% gelatin-coated dishes. Serum condition: Knockout DMEM (Life Technologies), 2 mM Glutamax (Gibco), 0.1 mM non-essential amino acids (Gibco), 15% ESC-grade fetal bovine serum (FBS) (Gibco), 0.1 mM β-mercaptoethanol, and leukemia inhibitory factor (LIF) (Millipore), 2i/Serum condition: above medium supplemented with 2i; 1 μM MEK inhibitor PD0325901 (Sigma) and 3 μM GSK3 inhibitor CHIR99021 (Sigma). 2i condition: serum free ESGRO Complete Basal medium (Millipore) with 0.1mM LIF and 2i as described above. For drug treatment, mESCs were grown in respective medium supplemented with 10 μM GSK-J4 (Sigma) for 96 h, 50 μM Zebularin for 96 h or 10 μM EPZ-6438 (BioVision) for 7 days. For all conditions cells were passaged in 48 h intervals, using accutase (Sigma) for detachment. Cell line was tested for mycoplasma contamination.

### Immunoblot Analysis

1×10^6^ cells were harvested for each growth condition, washed twice with phosphate buffered saline (PBS) and lysed in 500 μl of ice-cold radioimmunoprecipitation (RIPA) buffer (0.1% sodium deoxycholate, 0.1% SDS, 1% Triton X-100, 10 mM Hepes [pH 7.6], 1 mM EDTA, 5% glycerol and 140 mM NaCl) supplemented with Protease Inhibitor Cocktail (PIC, Roche) on ice for 10 min. For the drug treated conditions, 10 μl of lysate from the ChIP sample was used for the analysis directly. Lysates were homogenized by sonication for 8-10 cycles at high power, 30 seconds on/off in a Bioruptor sonicator (Cosmo Bio Co. Ltd.). Samples were boiled at 95 °C for 10 mins with 6×SDS sample buffer before loading onto 4–20% Tris-glycine gels (BioRad). Resolved proteins were transferred to nitrocellulose membranes using the Trans-Blot^®^ Turbo system (BioRad) according to manufacturer’s instructions. Membranes were then blocked for 1 h in 1% casein prepared in Tris-buffered saline and 0.1% Tween-20 (TBS-T) before blotting with respective primary antibodies diluted in TBST, overnight at 4°C. Blots were washed three times with TBST and incubated with secondary antibody in the same buffer for 1 h at room temperature (protect from light). Post three TBST washes, the membranes were imaged on a LI-COR Odyssey ^®^ FC system. Quantitation of signal and analysis was performed using the LI-COR Image studio software. Primary antibodies included total H3 1:10,000 (Active motif 39763), H3K4me3 1: 5000 (Millipore 04-745), H3K27me3 1: 5000 (Millipore 07-449). The secondary antibodies were IRDye^®^ 680RD anti-rabbit and IRDye^®^ 800CW anti-mouse (LI-COR) at 1:5000 dilution. Full western blot images have been included in the Supplementary Information.

### Mass Spectrometry

5×10^6^ mESCs per growth condition (serum or 2i) were harvested, washed once with PBS, spun down at 800 g for 5 min. The cell pellets were flash frozen and sent to ActiveMotif for their Mod Spec^®^ service. Briefly, histones were acid extracted, derivatized via propionylation, digested with trypsin, newly formed N-termini were propionylated, and then measured three seperate times using the Thermo Scientific TSQ Quantum Ultra mass spectrometer coupled with an UltiMate 3000 Dionex nano-liquid chromatography system. The data was quantified using Skyline, and represents the percent of each modification within the total pool of that tryptic peptide.

### MINUTE-ChIP

#### One-pot chromatin fragmentation and barcoding

MINUTE-ChIP and library preparation protocol is based on the Mint-ChIP protocol developed by the Bernstein lab ^12^, with modifications as follows: 1×10^6^ cells were harvested for each growth condition, washed twice with PBS and cell pellets were flash frozen at −80°C prior to use. Cells were resuspended in 50 μl PBS, lysed and digested to mono- to tri-nucleosomes fragments by adding 50 μl of 2x Lysis buffer (*100 mM Tris-HCL* [*pH 8.0*], *0.2% Triton X-100, 0.1% sodium deoxycholate, 10 mM CaCl_2_ and 1x PIC*) containing 2U/μl of micrococcal nuclease (New England BioLabs, M0247S) and incubating on ice for 20 min and then 37°C for 10 min. For each sample, 40 μl of the whole cell lysate containing the digested chromatin was taken forward into an overnight blunt end ligation reaction (End-It DNA repair kit and Fast-Link DNA ligation kit, Epicentre) with double stranded DNA adapters at 16°C. As in the original Mint-ChIP design, adaptors carried a partial SBS3 for Illumina sequencing flanked by a T7 RNA Polymerase for linear amplification. Between the SBS3 sequence and a 8bp sample barcode at the 3’ end, a 6bp randomized sequence was introduced, serving as a unique molecular identifier (UMI) (Figure 1a). UMI and sample barcode are ligated 5’ to the chromatin fragment and constitute the first 14 bases of read 1. The 4096 possible UMIs provide sufficient diversity to distinguish if two reads mapping to the exact same genomic location arose from a PCR amplification artifact or are indeed unique molecule. The adapter concentration was optimized to 2.5 μM / reaction to reduce adapter dimers. The ligation reaction was terminated with a lysis dilution buffer *(50 mM Tris-HCl* [*pH 8.0*], *150 mM NaCl, 1% Triton X-100, 50 mM EGTA, 50 mM EDTA, 0.1% sodium deoxycholate and 1x PIC)* and barcoded samples were combined into a single pool, spun down at 24,000 r.p.m. for 10 min at 4°C.

#### Immunoprecipitation

50μl Protein A/G magnetic beads (BioRad) were washed twice with PBS-T (PBS+ 0.1% Tween 20) and coupled to one of the following antibodies in the same buffer for 1 hr at room temperature with rotation: 3 μl H3 (Active motif 39763), 5 μl H3K4me3 (Millipore 04-745), 5 μl H3K27me1 (Cell signaling 5326S), or 5 μl H3K27me3 (Cell signaling 9733 or Diagenode C15410195 or Millipore 07-449). Beads were then washed quickly with RIPA buffer. 200-400 μl of the cleared lysate pool was added to the pre-coupled magnetic beads and parallel ChIP assays were incubated further for 4 h at 4°C with rotation. 5% of the above volume was saved as the input pool and processed through the remaining protocol in a manner similar to the IPs. Next, the beads were washed (RIPA, RIPA high salt, LiCl and TE buffer) resuspended in ChIP elution *buffer (10 mM Tris-HCl* [*pH 8.0*], *1 mM EDTA, 0.1% SDS, and 300 mM NaCl)* containing 0.25 mg/mL Proteinase K and eluted at 63°C for 1 h.

#### Linear amplification and library preparation

The native ChIP DNA (fragments longer than 100bp) was isolated using 1x SPRI beads (Beckman Coulter) and set up in an overnight *in vitro* transcription reaction (HiScribe T7 Quick High Yield RNA Synthesis kit, New England BioLabs). The resulting RNA was purified using Silane beads (Life Technologies) and an **RNA 3’ adapter** was ligated onto it using T4 RNA ligase, truncated (New England BioLabs) for 1 h at 25°C. The mixture was subsequently supplemented with components of the reverse transcription reaction (SuperScript III First-Strand Synthesis SuperMix, Life Technologies) to produce cDNA, primed using the ligated 3’ adapter. Final libraries for each ChIP were produced using 150-200 ng of purified cDNA in a PCR reaction (High-Fidelity 2x master mix, New England BioLabs) for 8 cycles with 0.2 μM primers that carried a second 8bp barcode sequence. Quality assessment and concentration estimation of the purified DNA was done using the Qubit (Life Technologies) and BioAnalyzer (Agilent). Each library was then diluted to 4 nM and combined into a single pool before sequencing on the Illumina NextSeq500 platform.

### MINUTE-ChIP Quantitative Mapping Pipeline

#### Demultiplexing and deduplication

Sequencing was performed using 50:8:34 cycles (Read1:Index1:Read2) Illumina bcl2fastq was used to demultiplex paired-end sequencing reads by 8nt index1 read (PCR barcode). NextSeq lanes were merged into single fastq files, creating the primary fastq files. Read1 starts with 6nt UMI and 8nt barcode in the format NNNNNNABCDEFGH. A sample sheet was used with four columns corresponding to sample name, replicate name, barcode sequence, primary fastq name. For each line in the sample sheet, reads matching the barcode sequence were extracted from the primary fastq file, allowing up to two mismatches. Duplicate reads identified by identical first 24nt of read1 were discarded. Read pairs matching parts of the adaptor sequence (SBS3 or T7 promoter) in either read1 or read2 were removed. Demultiplexed and deduplicated reads were written into sample-specific fastq file used for subsequent mapping.

#### Mapping

Sample-specific paired fastq files were mapped using botwie2 (v2.3.4.3) using --fast parameter to the mouse genome (mm9). Alignments were processed into sorted BAM files with samtools (v1.9). Blacklisted regions were removed from BAM file using BEDTools (v2.27.1). Reads were also mapped to a metagenome of RepBase murine repetitive sequences using botwie2.

#### Generation of coverage tracks and quantitative scaling

Input coverage tracks with 1bp resolution in BigWig format were generated from BAM files using deepTools (v3.1.0) bamCoverage and scaled to a reads-per-genome-coverage of one (1xRPGC, also referred to as ‘1x normalization’) using mm9 genome size 2654895218. ChIP coverage tracks were generated from BAM files using deepTools (v3.1.0) bamCoverage. Quantitative scaling amongst samples within each pool was based on their total mapped read counts, as well as the total mapped read counts of the respective input data. One condition in the pool served as a reference (Serum condition, no treatment), which was scaled to 1xRPGC. All other samples were scaled relative to the reference using an explicit scaling factor. The RPGC was calculate by the following formula: (#mapped[ChIP] / #mapped[Input]) / (#mapped[ChIP_Reference] / #mapped[Input_Reference]).

#### Genome statistics

Total mapped read counts from BAM files were used calculate relative global levels of histone modifications. Statistics on genomic bins or custom intervals were calculated from scaled BigWig files using bwtool summary. For calculation of enrichments in repetitive regions, mapped read counts for ChIP and Input of the repetitive sequences were Put in relation to the respective total mapped reads counts of the mm9 genome and scaled with the same scaling factor.

#### Track visualization

Heatmaps were generated from scaled BigWig files and clustered with SeqPlots using a 50bp moving window average smoothing. Density plots were generated directly from BAM files using ngs.plot with modifications: the in-built genome averaging was removed and replaced by a scaling factor calculated from the aligned read counts as above (#mapped[ChIP] / #mapped[Input]) / (#mapped[ChIP_Reference] / #mapped[Input_Reference]). Equivalent plots can be generated using SeqPlots from scaled BigWig files. For IGV visualization, a variable smoothing was applied depending on the size of the genomic region.

#### Quality control

Picard (v2.10.3) was used to determine insert size distribution, duplication rate, estimated library size. For the latter, primary reads were demultiplexed separately from the procedure described above retaining all duplicate reads and mapped using bowtie2 before running Picard MarkDuplicates. For estimation of cross contamination, mm9 and hg19 fasta files were combined to build a hybrid botwie2 index and used for mapping the reads.

#### GEO datasets

Following published datasets were used: RNA-Seq Serum vs 2i ^3, 26^ (GSM590124, GSM590125, GSM590126, GSM590127, GSM2345133, GSM2345139), CpG methylation ^7^ (GSM1127953, GSM1127954), H3K37me3 ^3, 27^ (GSM590114, GSM590113, GSM2229356, GSM2229359, GSM2229364)

## Data availability

For each sample demultiplexed and deduplicated reads were submitted to the GEO under GSE126252. Additional genomic files, pseudocode and supplementary data files are available on Mendeley Data: http://dx.doi.org/10.17632/s23bhg4xjv.1

## SUPPLEMENTARY MATERIAL

### Supplementary Figure Legends

**Supplementary Figure 1 – related to Figure 1.**
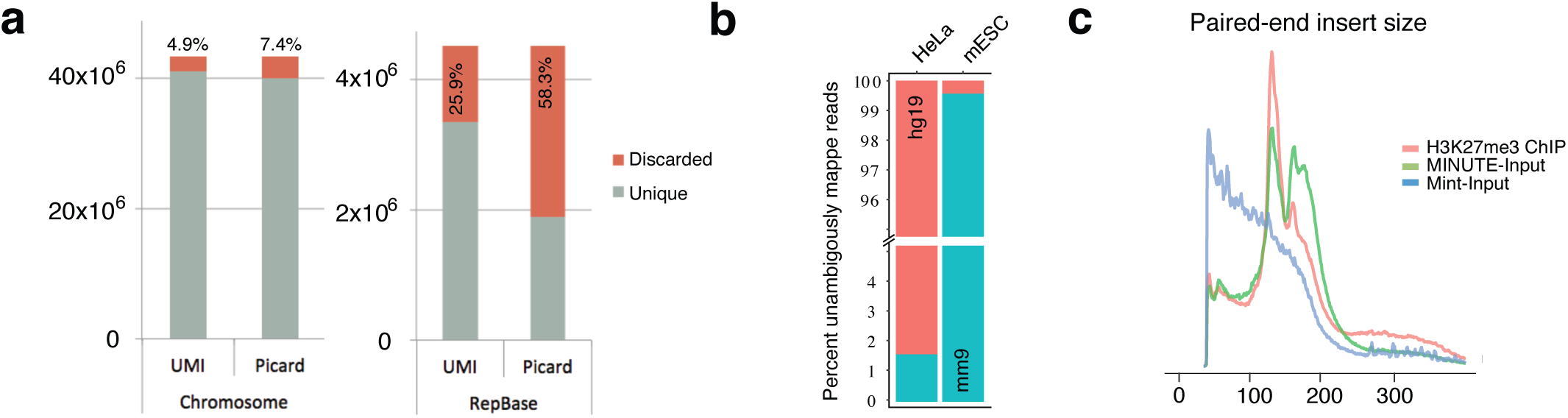
**a** Deduplication using unique molecular identifier. Total mapped read counts of the Input (Pool1) before and after demultiplexing with Picard (http://broadinstitute.github.io/picard/, relies on location of read1 one and read2) or with JE ^28^ using the UMI information. Reads were mapped to mm9 genome (left) and a metagenome with all murine repetitive sequences from RepBase ^29^ (right). **b** Analysis of barcode cross-contamination using a human HeLa cell sample in Pool1. One HeLa and one mESC sample were aligned to a hg19/mm9 metagenome and only uniquely mapped reads to either human or mouse chromosomes were counted. **c** Fragment insert size distribution determined from aligned BAM file, comparing the input library prepared using RA3 cDNA primer (see Figure 1a, MINUTE-ChIP) or random hexanucleotide priming according to original Mint-ChIP protocol.

**Supplementary Figure 2 – related to Figure 1.**
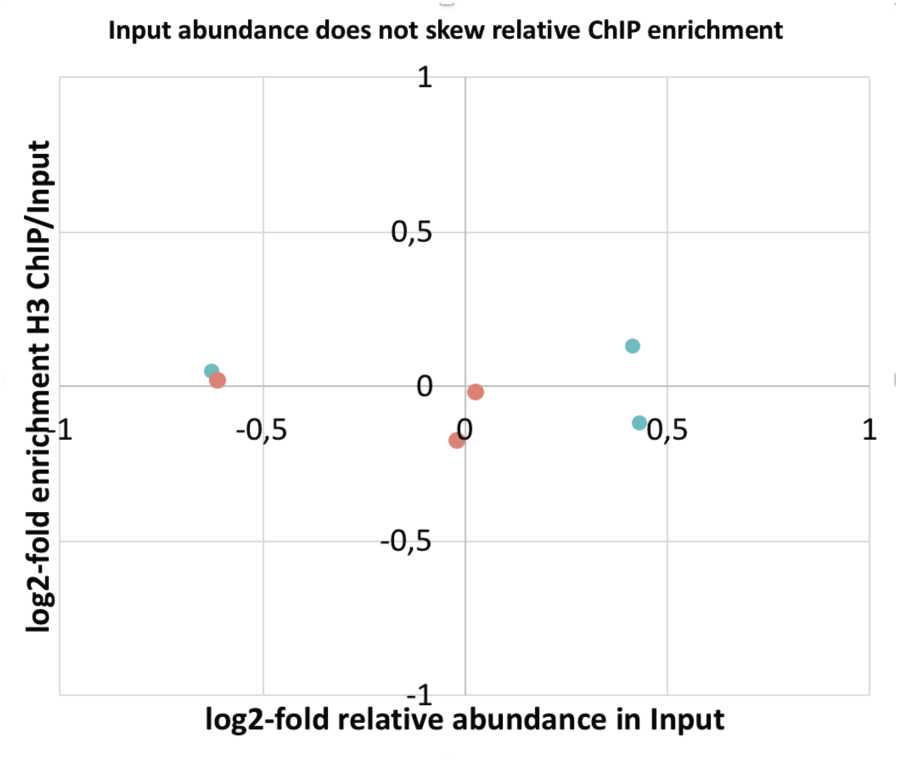
Plot comparing the original abundance (log2) of sample barcodes in Pool1 with their fold-enrichment (log2) after an H3 ChIP. As expected there is no correlation between the two variables, e.g. low abundant barcodes are neither more nor less biased to become enriched during IP.

**Supplementary Figure 3 – related to Figure 1.**
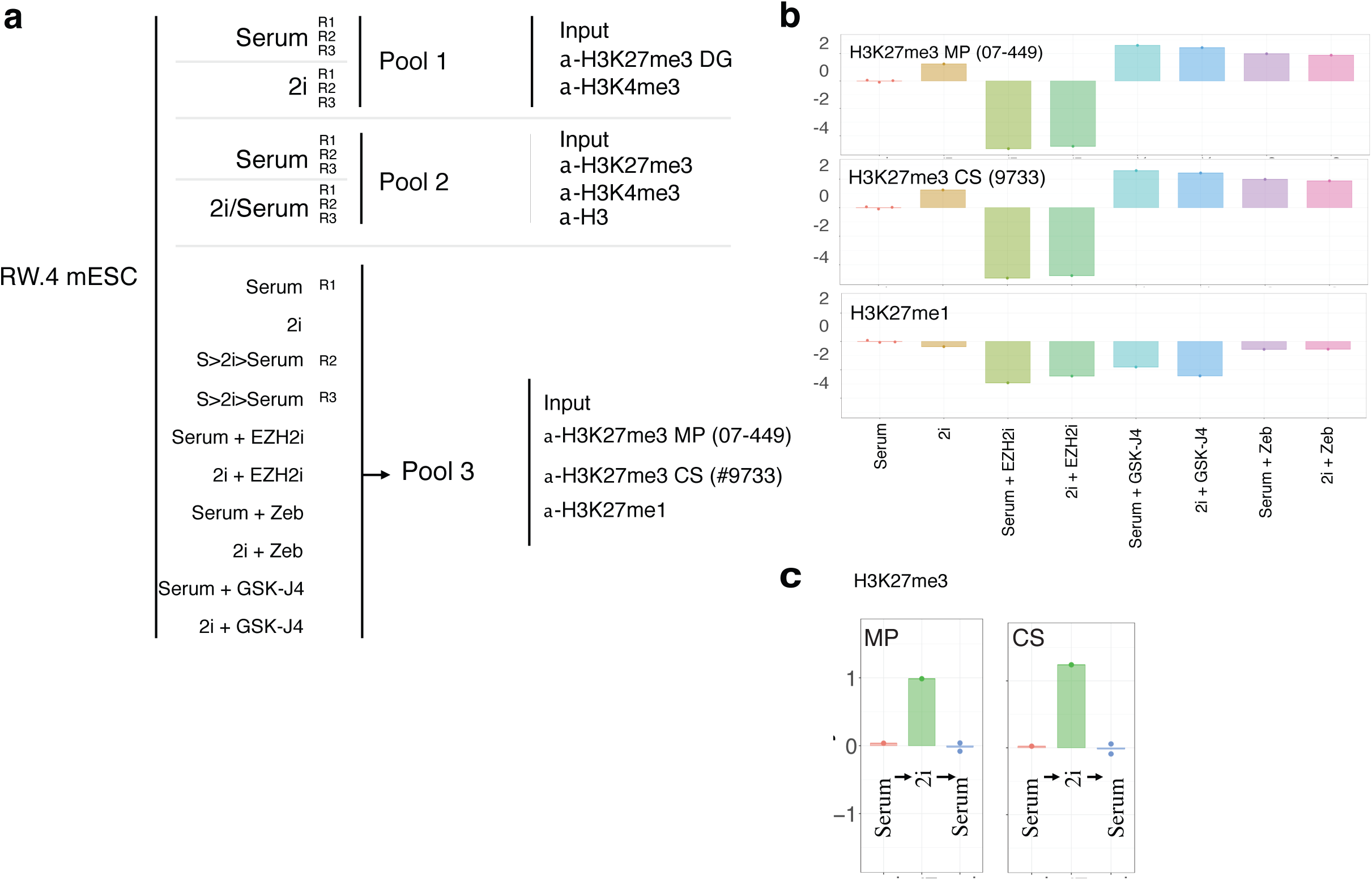
**a** Overview of the three MINUTE-Pools used and ChIPs performed from each pool in the present study. Serum is abbreviated with “SL” here **b** Quantitative comparison of H3K27me3 and H3K27me1 from total mapped read counts after ChIP compared to input abundance. Full panel relating to Figure 1f. **c** Two of the three Serum samples in Pool3 were shifted to 2i and back for three passages. Global H3K27me3 returns to pre-2i levels.

**Supplementary Figure 4 – related to Figure 1.**
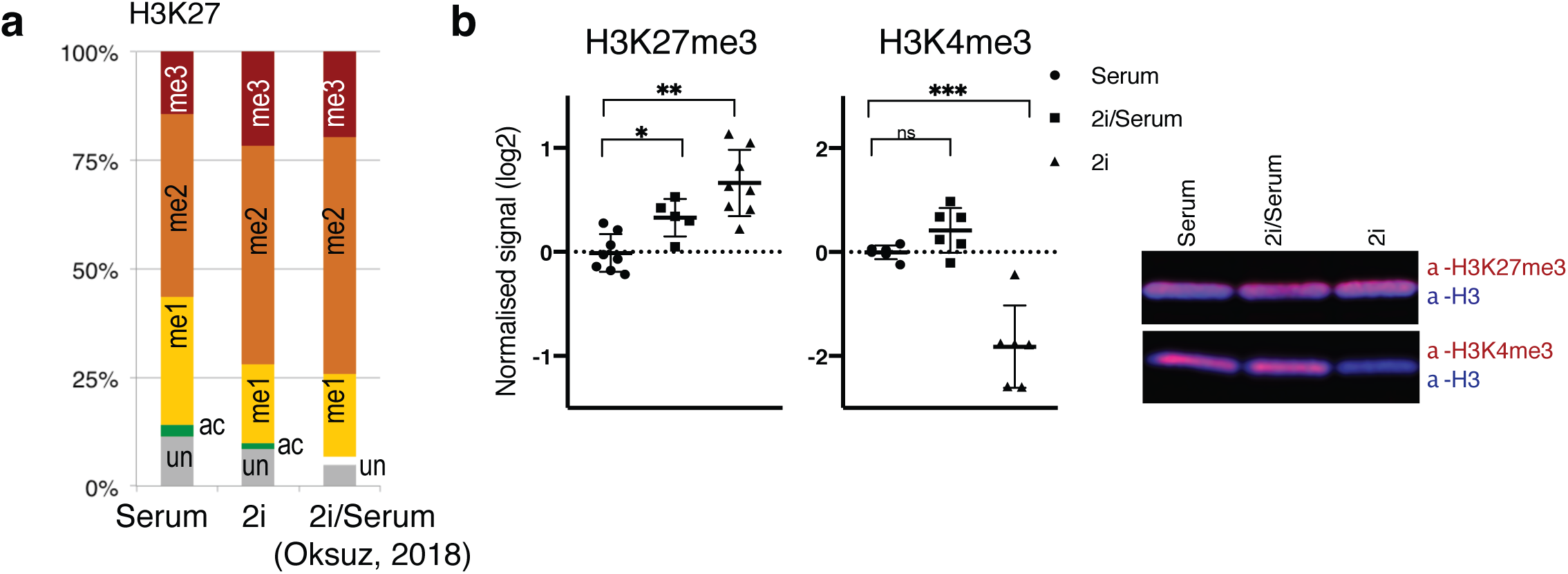
Validation of quantitative differences between Serum and 2i condition using western blot and mass spectrometry. a relative quantitative assessment of H3K27me3 levels using mass spectrometry (ModSpec, ActiveMotif) in Serum and 2i (one sample each). K27me3 on histone H3.1 is estimated to be 14% in serum and 22% in 2i conditions. A recent study performed a similar quantitation in 2i/Serum conditions (right) and reported 20% ^30^. H3K27ac was not reported in the latter study. Note that H3K4me3 does not produce adequate peptides for quantification with this protocol. **b** Quantitation of H3K27me3 and H3K4me3 using two-color IR western blot, in Serum, 2i and 2i/Serum conditions. T-test was performed on triplicate samples (* = p<0.05; ** = p<0.005). Representative image of the replicates is shown.

**Supplementary Figure 5 – related to Figure 1.**
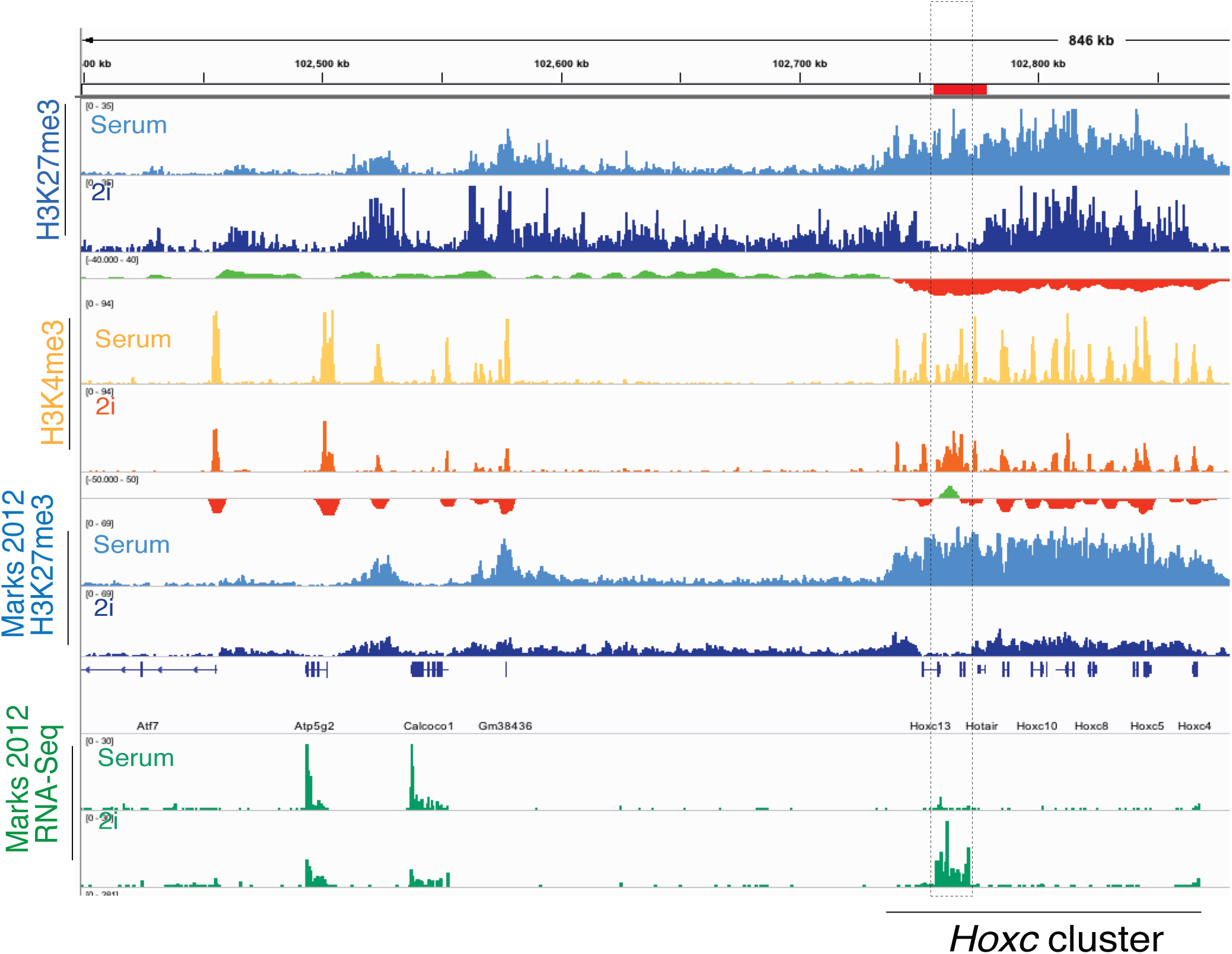
Comparison of data from ours study with published H3K27me3 dataset ^3^. Qualitative similarities and the difference in global scaling introduced by per-sample internal normalization are apparent. Nevertheless. previously reported region in *Hoxc* cluster shows dramatic loss of H3K27me3 and transcriptional induction, as reported ^3^, also a concomitant gain in H3K4me3.

**Supplementary Figure 6 – related to Figure 1.**
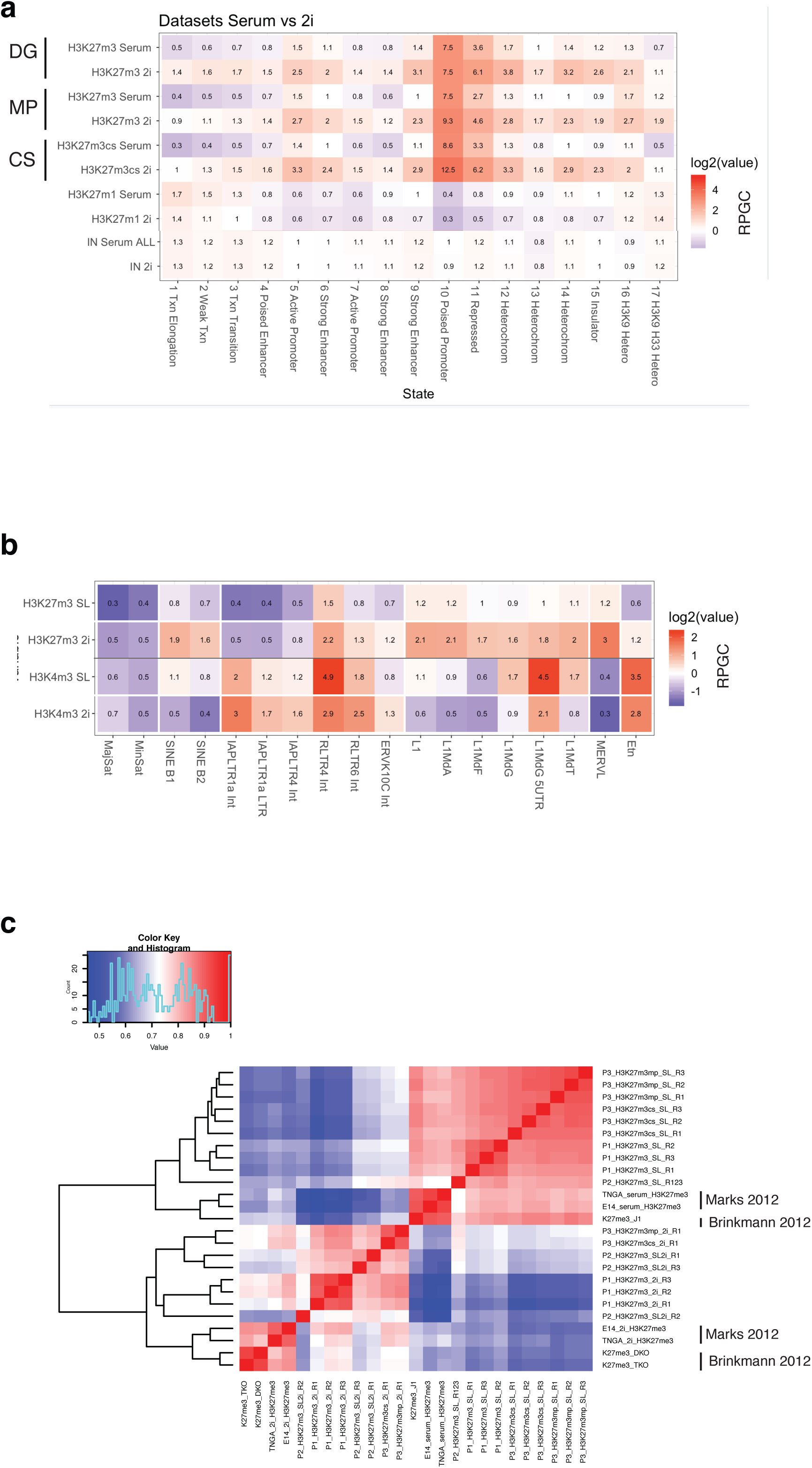
**a** Comparison of the three H3K27me3 datasets comparing Serum and 2i conditions acquired from two independent pools and three different H3K27me3 antibodies. Relative enrichments of H3K27me3 in Serum and 2i conditions, over 17 functionally defined chromatin states (ChromHMM). **b** Relative enrichments of H3K27me3 and H3K4me3, in Serum and 2i conditions, over repetitive regions of the genome using reads mapped to a metagenome of repetitive sequences from RepBase ^29^. **c** Hierarchical clustering shows that present datasets cluster with prior H3K27me3 data from Serum vs 2i comparison ^3^. Pearsson correlation was calculated on 10kb bins of the mm9 genome after removing outliers and blacklisted regions.

**Supplementary Figure 7 – related to Figure 2.**
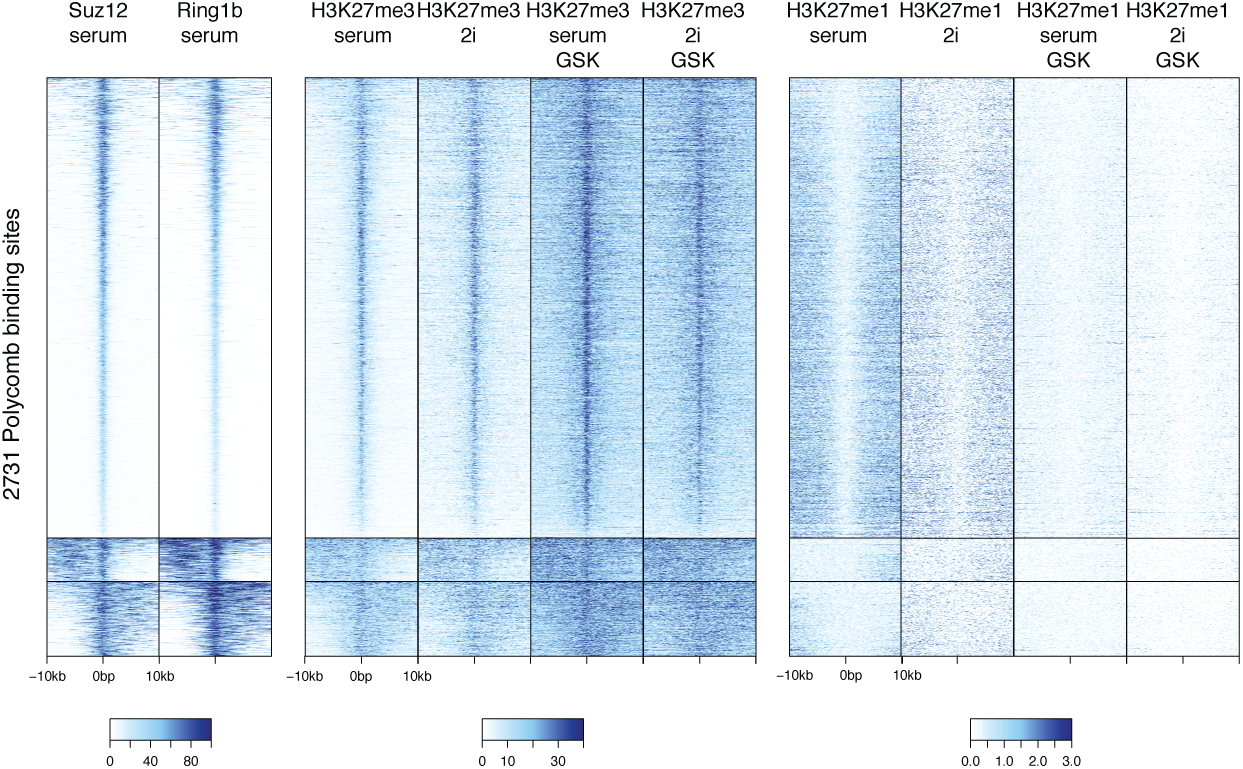
Heatmaps over 2731 PcG target sites defined by EZH2 or Ring1b binding. Peak locations were from acquired from published datasets (USCS Genome Browser and ^4^). Suz12 and Ring1b ^4^ were plotted and used for clustering alongside of H3K27me3 and H3K27me1 data.

**Supplementary Figure 8 – related to Figure 2.**
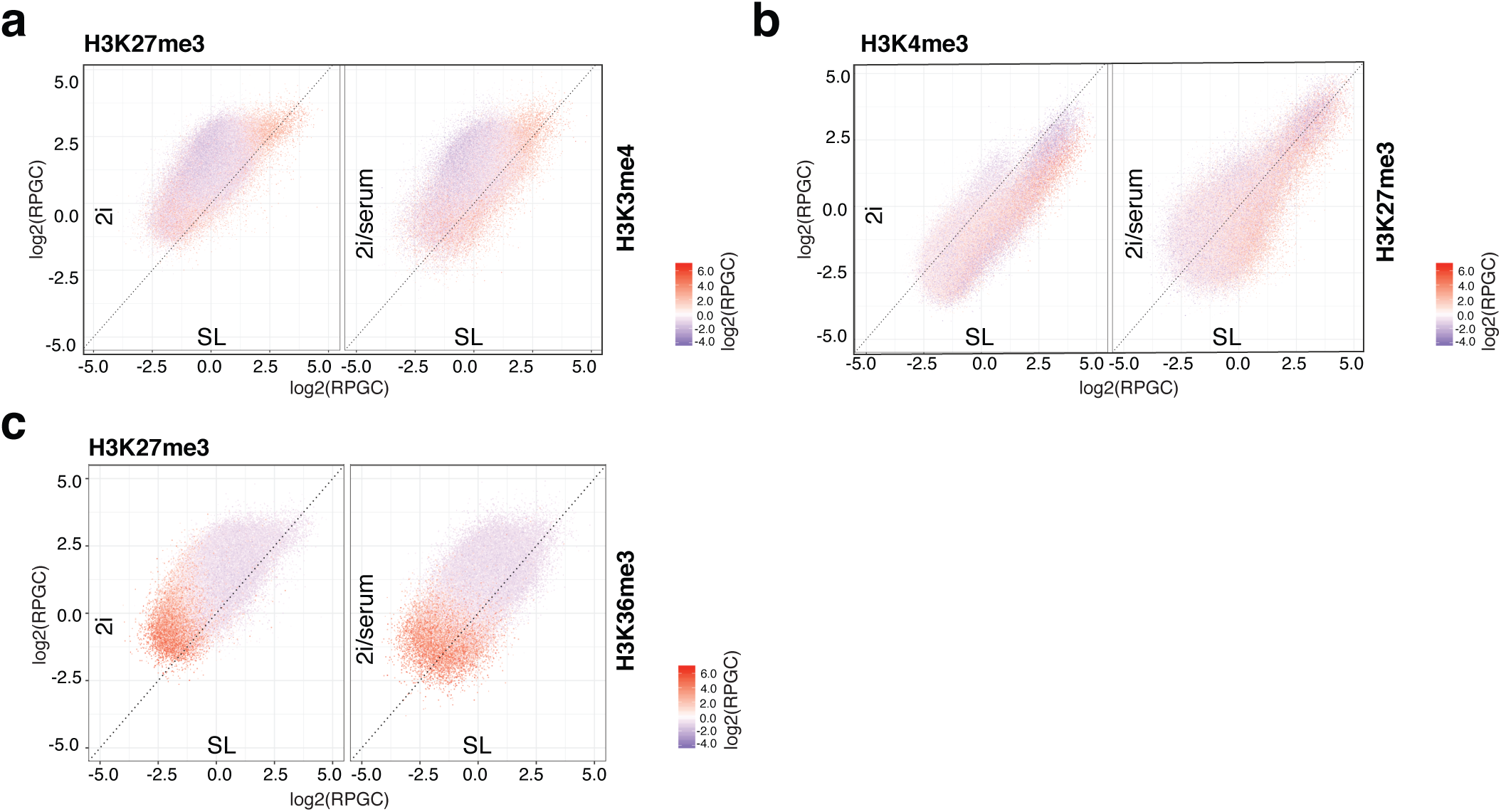
**a** Genome-wide analysis of 10kb windows. Levels of H3K27me3 (expressed as RPGC relative to Serum) were determined per bin and plotted in a 2D comparison. A color scale was used to overlay H3K4me3 levels of each bin. H3K27me3 increased almost universally across the genome in 2i/serum or 2i. As an exception, the most extreme high regions of H3K27me3 (which coincided with high H3K4me3) did not further increase in H3K27me3. **b** Levels of H3K4me3 (expressed as RPGC relative to Serum) were determined per bin and plotted in a 2D comparison. A color scale was used to overlay H3K27me3 levels of each bin. H3K4me3 levels on the other hand were globally unchanged in 2i/serum. Loss of H3K4me3 in 2i was limited to H3K4me3-high regions, corresponding to active (H3K27me3 low) and bivalent (H3K27me3 high) regions. **c** H3K27me3 bins as in a) were overlaid with H3K36me3 density color, showing that H3K36me3-high regions (regions of highly active transcriptional elongation) were most depleted in H3K27me3, but still gained H3K27me3 in 2i condition.

**Supplementary Figure 9 – related to Figure 2.**
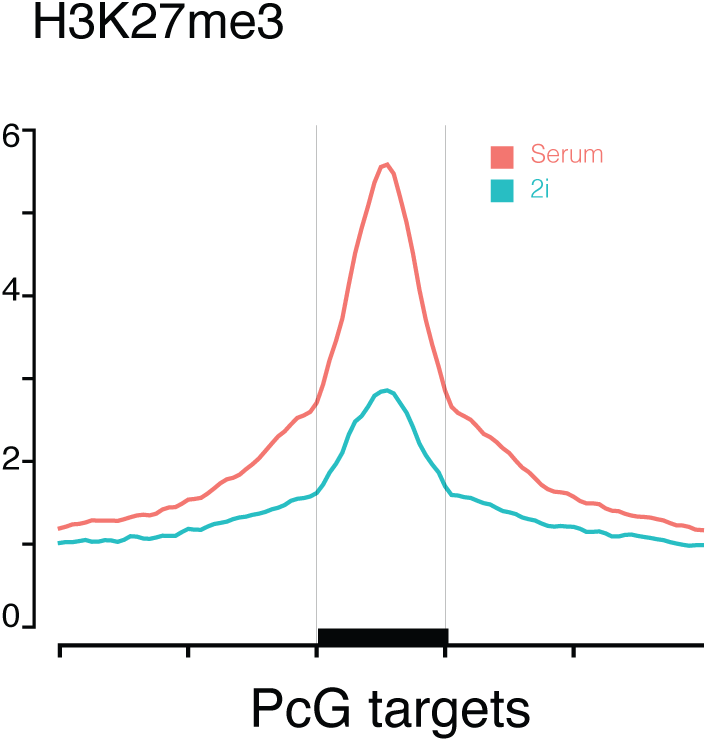
Average plot over 2731 PcG target sites using per-sample normalization. Because of increase in background H3K27me3 levels, the peak size appears dramatically reduced.

**Supplementary Figure 10 – related to Figure 2.**
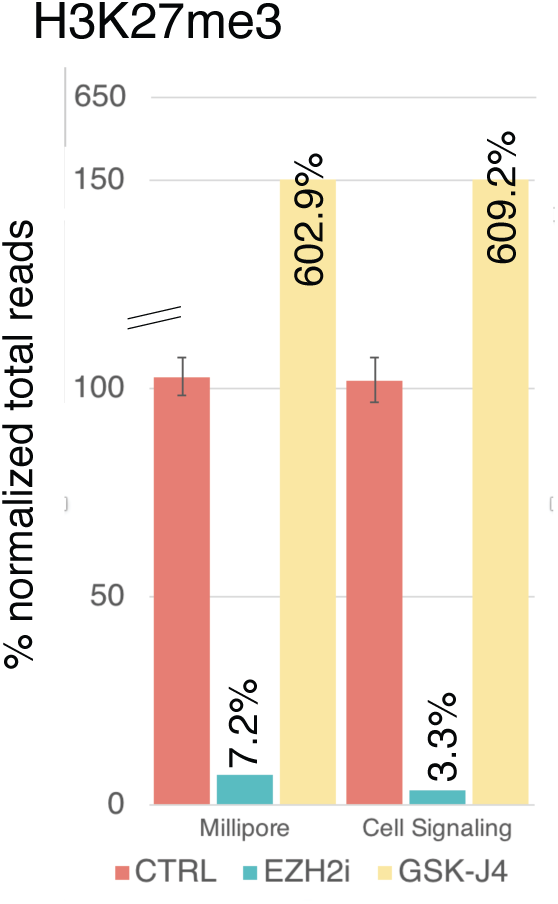
Background and dynamic range of MINUTE-ChIP method as determined by modulating H3K27me3 levels. Relative abundance of barcodes from Pool3 corresponding to Serum (CTRL), EZH2i and GSK-J4 inhibitor treatments in Serum.

**Supplementary Figure 11 – related to Figure 2.**
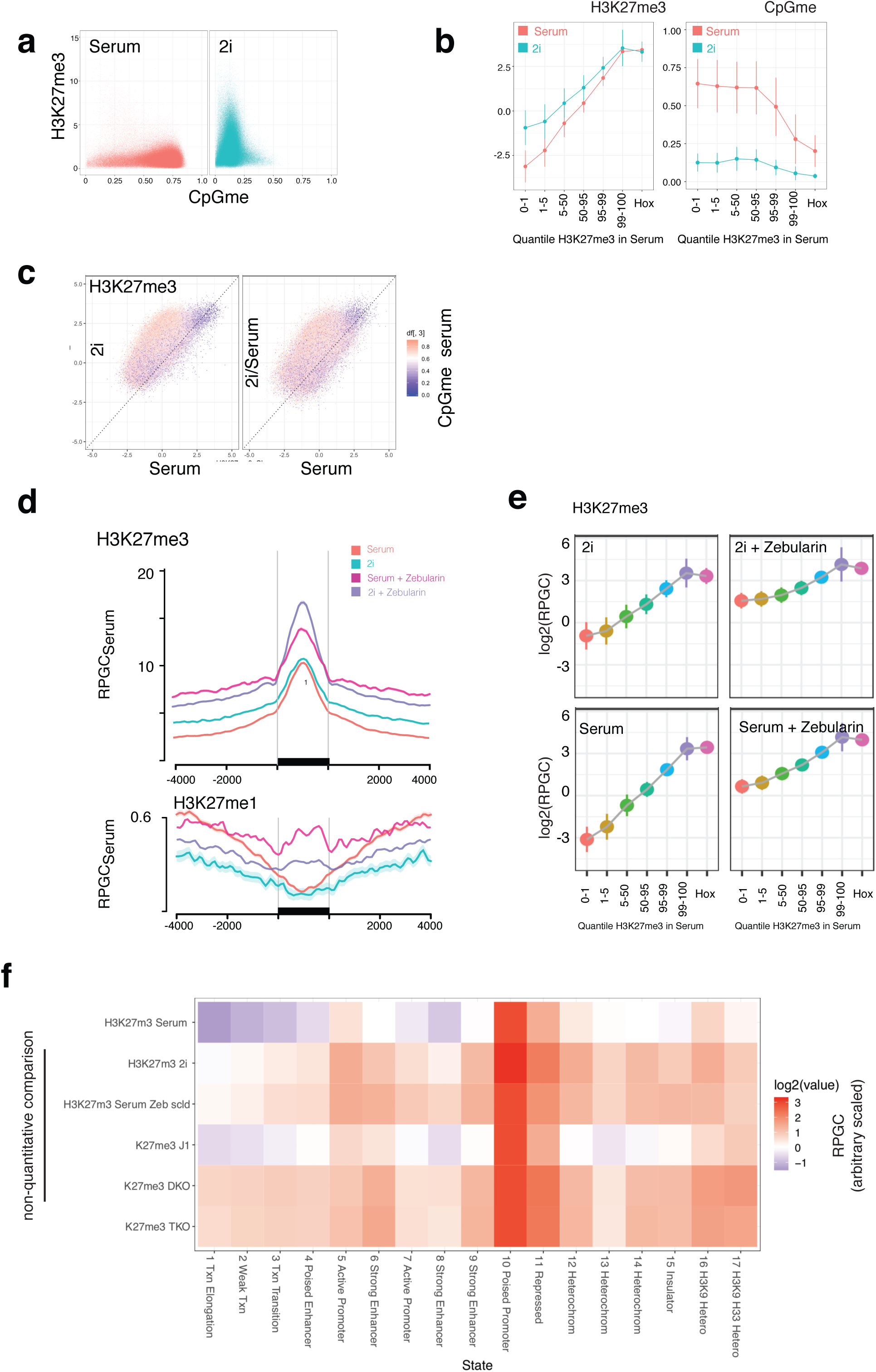
H3K27me3 follows loss of CpG methylation. **a** H3K27me3 and CpG methylation ^7^ levels in Serum and 2i conditions were determined in 10kb windows and plotted against each other. The gain of H3K27me3 observed in the bulk of 10kb windows in 2i is concomitant to a loss of CpG methylation. **b** 10kb bins were partitioned in 1, 5, 50, 95, 99th quantiles according to H3K27me3 levels in Serum. Regions overlapping with hox genes were analyzed separately. Overlay of their levels in 2i shows an increase inversely related to the original level, with the most devoid regions showing the largest fold-change. CpG methylation and H3K27me3 was largely anticorrelated **c** Levels of H3K27me3 (expressed as RPGC relative to Serum) were plotted in a 2D comparison. A color scale was used to overlay the CpGme levels of each bin. Bins high in H3K27me3 were constitutively unmethylated and did not gain H3K27me3. **d** Average profile of H3K27me3 and H3K27me1 levels at PcG target, in Serum and 2i condition, as well as after treatment with DNMT inhibitor Zebularin for 4 days. Y axis is normalized to RPGC in Serum. **e** Quantile analysis as in b) shoes that Zebularin treatment promotes hypomethylation across all quantiles, and a particularly strong increase in H3K27me3 at the low end. **f** Comparison of Zebularin treatment and DNMT3a/b knockout (DKO), DNMT1/3a/3b knockout (TKO), in modulating H3K27me3 levels accross functional chromatin states. Since available data for DNMT KO was not performed quantitative, we normalized data according to State 10. Zebularin treatment and DNMT DKO/TKO exhibit similar shifts in H3K27me3, which is qualitatively similar but not identical to 2i. Note that DNMT DKO and TKO profiles also clustered with 2i samples in a correlation analysis (Supplementary Figure 6c)

**Supplementary Figure 12 – related to Figure 2.**
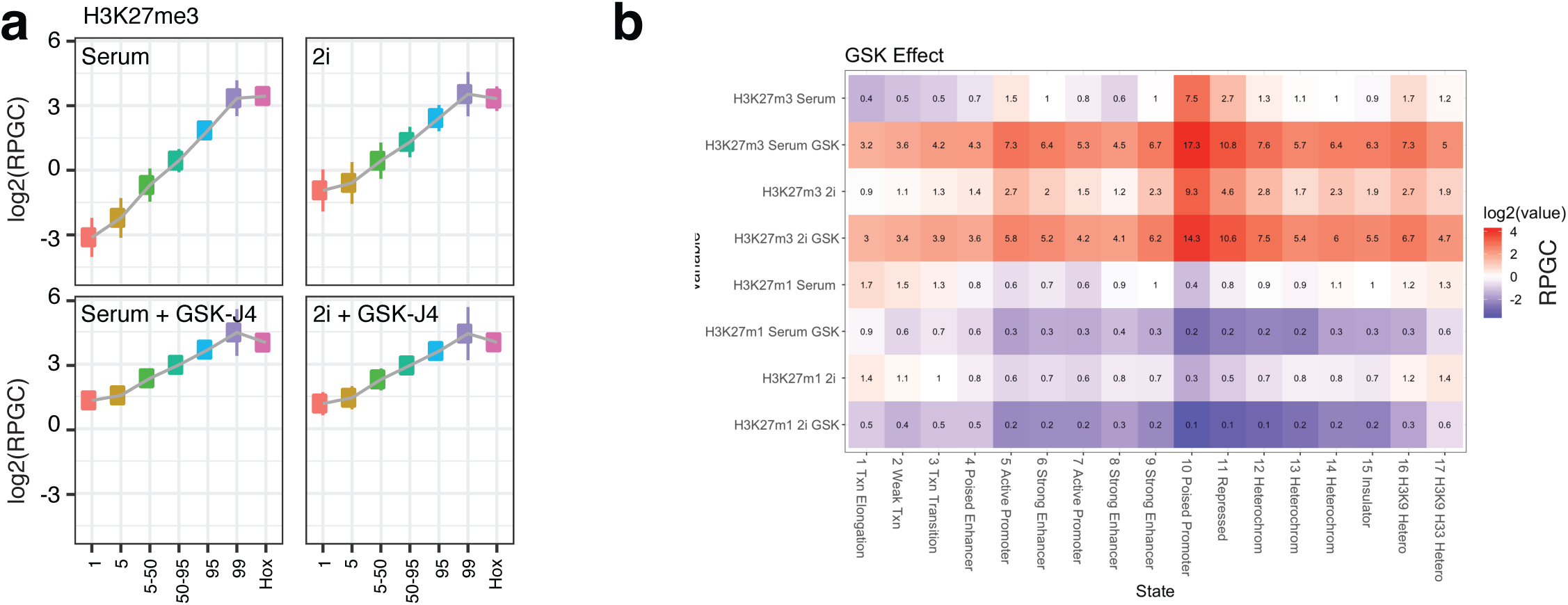
Inhibiting demethylases globally increases H3K27me3 levels. **a** 10kb bins were partitioned in 1, 5, 50, 95, 99th quantiles according to H3K27me3 levels in Serum. Regions overlapping with hox genes were analyzed separately. Overlay of the GSK-J4 treatment (4 days) shows that H3K27me3 levels are increased across all quantiles. **b** GSK-J4 effect on H3K27me3 and H3K27me1 levels analyzed by functional chromatin states. Increase in H3K27me3 upon GSK-J4 treatment is mirrored by a reduction of H3K27me1, the product of H3K27me3 demethylases.

**Supplementary Figure 13 – related to Figure 3.**
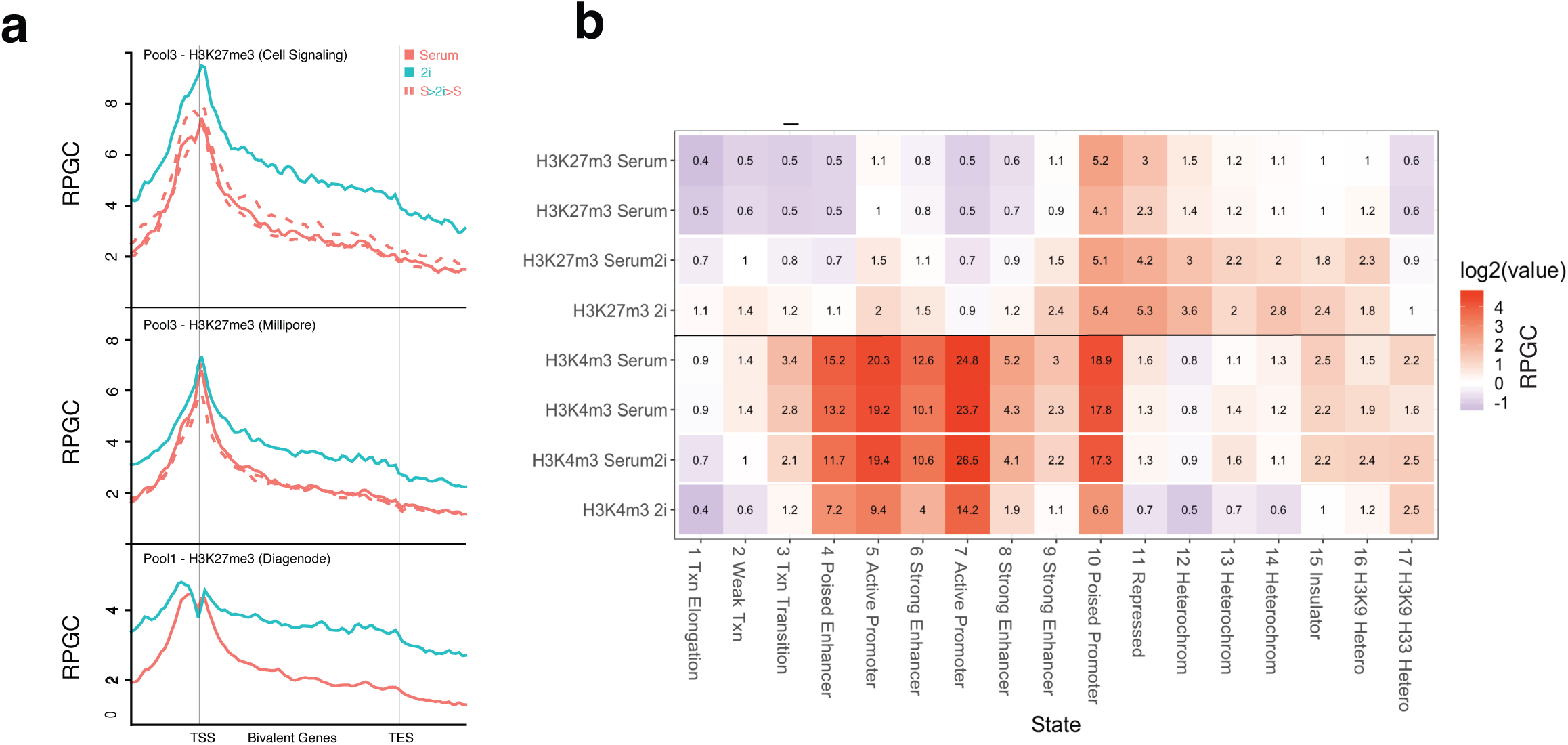
**a** Average plots of H3K27me3 levels at bivalent genes produced by three different antibodies from 2 different pools. **b** H3K27me3 and H3K4me3 levels in Serum, 2i, 2i/Serum by functional chromatin state. Also see global levels shown in Figure 1c. It is apparent that changes to the H3K27me3 landscape in 2i are largely mirrored by 2i addition to Serum condition (top). In contrast, 2i/Serum is very similar to both Serum datasets, and does not exhibit the strong reduction of H3K4me3 in 2i. H3K4me3 is lost most dramatically (>2.5-fold reduction) from State 10 “Poised_Promoters” that are bivalent. Active promoters are reduced ∼2-fold.

**Supplementary Figure 14 – related to Figure 3.**
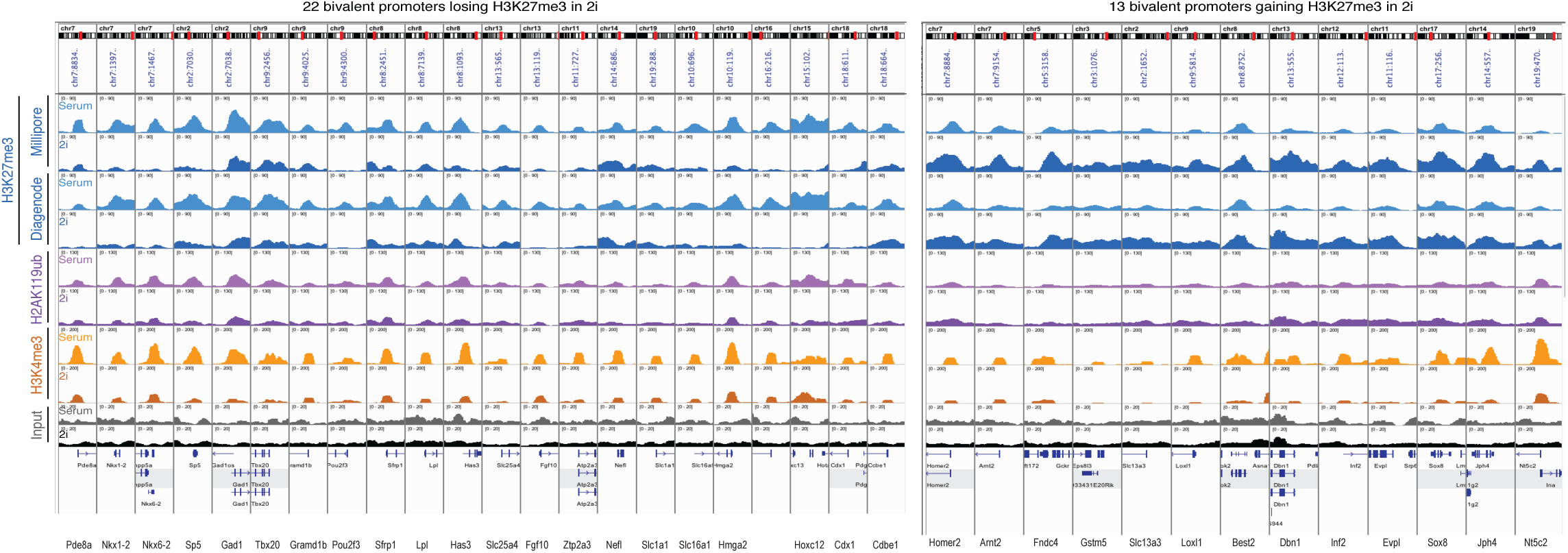
Examples of bivalent genes that lose or gain H3K27me3 in 2i condition more than two-fold. Notably, all promoters also lose H3K4me3, independent of the gain or loss of H3K27me3.

**Supplementary Figure 15 – related to Figure 3.**
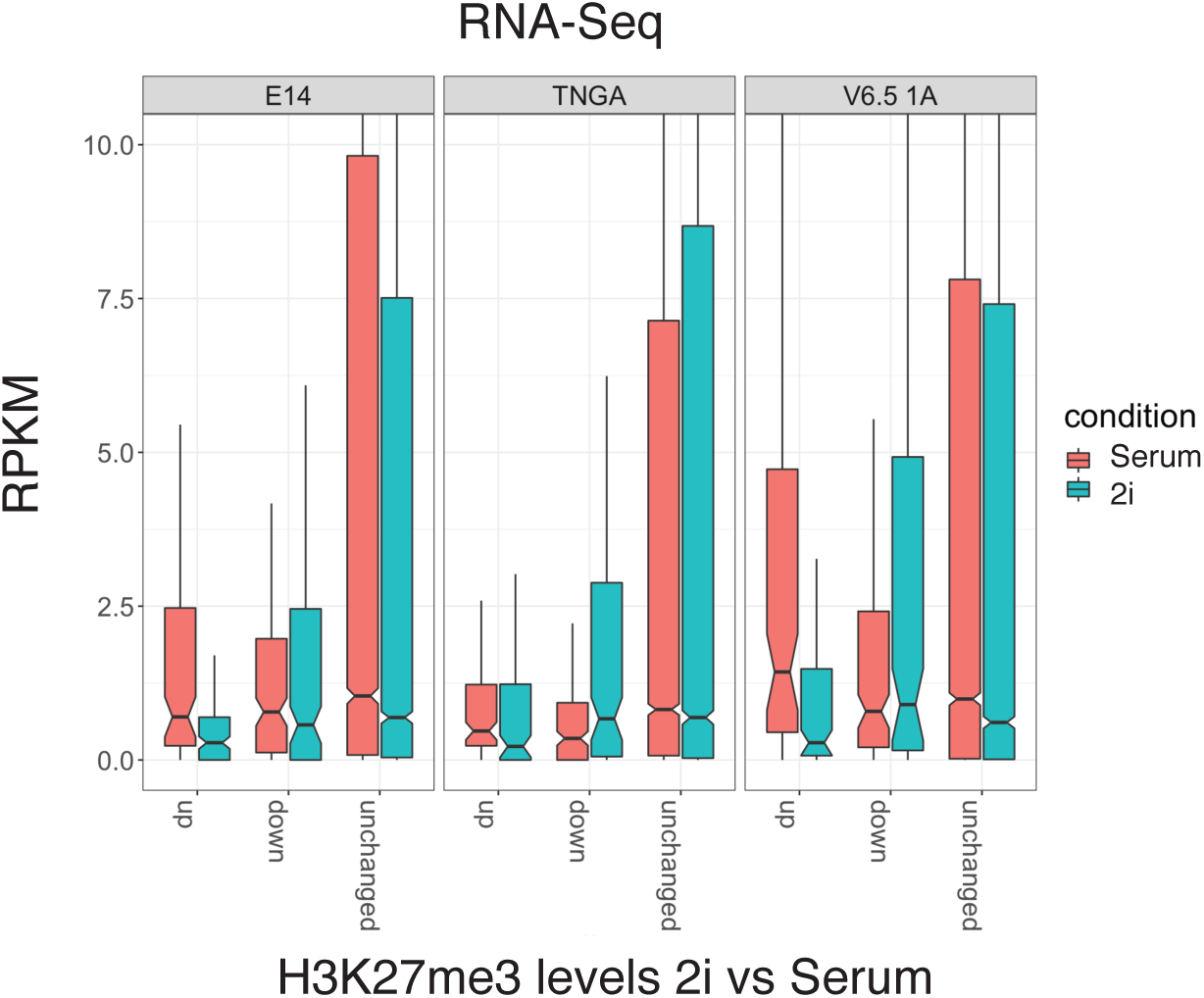
RNA-Seq expression of bivalent genes in Serum and 2i. Those genes that consistently gain (n=126) or lose (n=171) H3K27me3 were analyzed separately. RNA-Seq data from different studies was used ^3, 26^.

### Supplementary Tables

Attached in separate Excel file

Supplementary Table 1 - Sequences of oligos used for adaptors and PCR primers

Supplementary Table 2 - MINUTE-ChIP Pools and Quantification

